# Simultaneous Real-time Imaging of Neurofluid and Neurovascular Dynamics Using Ultrafast Flow-weighted Echo-Planar Imaging

**DOI:** 10.64898/2026.01.15.699612

**Authors:** Amir H. Shaker, Amir H. G. Arani, Bryn A. Martin, Aristeidis Sotiras, Cihat Eldeniz, Helia Hosseini, Joshua S. Shimony, David D. Limbrick, Jennifer M. Strahle, Arash Nazeri

## Abstract

Cerebrospinal fluid (CSF) circulation is tightly coupled to cerebral blood flow under the fixed-volume constraint of the cranial vault, with cardiac pulsations and respiration acting as dominant physiological drivers. Disruption of these flow dynamics has been implicated in various neurological disorders, motivating the need for imaging methods that capture CSF and vascular flow simultaneously and in real time. Conventional phase contrast MRI (PC-MRI) provides quantitative CSF velocity measurements but relies on cardiac gating and velocity encoding, which limit temporal resolution, dynamic range, and sensitivity to non-cardiac fluctuations. Here, we introduce self-gated ultrafast real-time flow-weighted echo-planar imaging (SURF-EPI), an inflow-weighted approach in which signal intensity reflects the replacement of saturated spins by freshly inflowing, unsaturated water molecules, yielding higher signal with higher flow. This allows simultaneous assessment of arterial inflow, venous outflow, and CSF motion at high temporal resolution. SURF-EPI was acquired at the C2–C3 spinal level, enabling real-time imaging of CSF flow dynamics in the cervical spinal canal alongside arterial and venous blood flow in major cervical vessels (frame-rate: 21.7 Hz). In addition to frequency-domain analyses, we leveraged intrinsic arterial signal fluctuations as a timing reference to reconstruct cardiac-resolved CSF dynamics for individual cardiac cycles without external physiological recordings. Frequency-domain analysis revealed distinct spectral signatures in CSF flow compared with neurovascular flow, including broadened cardiac peaks and enhanced respiratory modulation, particularly within the ventral spinal canal. In contrast, dorsal CSF showed increased power within the cardiac frequency band, higher coherence with cervical vasculature at the cardiac frequency, and reduced non-cardiac contributions. Time-domain analysis showed strong correlation between CSF flow waveforms derived from SURF-EPI and PC-MRI. Beyond ensemble-averaged waveforms, SURF-EPI enabled beat-to-beat analysis, revealing substantial cycle-to-cycle variability in CSF flow that is not captured by time-averaged gated approaches. Together, these findings establish SURF-EPI as a rapid, complementary framework to PC-MRI, enabling real-time neurofluid imaging with integrated time– and frequency-domain characterization of CSF and neurovascular flow dynamics.

## 1. Introduction

Cerebrospinal fluid (CSF) circulation plays a critical role in maintaining brain fluid homeostasis and clearing of metabolic waste^1,2^. Disruptions in this tightly regulated system have been implicated in various brain disorders ranging from hydrocephalus, idiopathic intracranial hypertension to stroke and neurodegenerative disorders^1,3,4^. The rigid cranial vault imposes a fixed-volume constraint (Monro–Kellie doctrine), such that CSF flow dynamics are inseparably linked to cerebral blood flow^5,6^. Within this confined system, cardiac pulsations and respiratory cycles are the primary drivers of CSF and cerebral blood flow^7,8^. Accordingly, capturing these interactions in real time through simultaneous imaging of vascular and CSF flow is critical for understanding neurofluid physiology in health and its pathological disruption in disease.

Cardiac-gated phase-contrast (PC) MRI has long served as the standard imaging approach to quantify CSF velocity and flow across the cardiac cycle^9,10^. However, reliance of conventional PC-MRI on cardiac gating restricts measurements to a time-averaged representation, overlooking the real-time fluctuations driven by respiration and other non-cardiac factors^11,12^. Real-time PC MRI, by contrast, circumvents the need for gating and allows dynamic flow assessment^12–14^, enabling evaluation of respiratory and other non-cardiac drivers of CSF flow, as well as characterization of flow variability across cardiac cycles. Nonetheless, the dynamic range of both conventional and real-time PC methods are constrained by velocity encoding (VENC) setting, which sets the maximum measurable velocity without aliasing^10^. While lowering VENC thresholds can enhance phase signal-to-noise ratio (SNR) for detecting slow flow, it also increases the risk of velocity aliasing^15^. More fundamentally, the velocity constraints imposed by VENC preclude simultaneous measurement of neurovascular blood and CSF flow dynamics.

To overcome these limitations, alternative imaging strategies are being explored that allow simple CSF flow imaging using fast imaging methods^7,16^. Inflow-weighted approaches exploit flow-related signal enhancement. Freshly entering, unexcited CSF produces high signal intensity, whereas stationary fluid signal within the slab remains suppressed because its long T1 prevents appreciable recovery of longitudinal magnetization at short repetition times (TRs)^7,17^. Previous studies employing whole-brain echo-planar imaging (EPI) have successfully mapped unidirectional CSF flow in a few edge slices, restricted to flow entering the imaging slab. However, these methods require extended acquisition times and offer relatively low temporal resolution (250 – 500 ms)^16–20^.

Here, we introduce self-gated ultrafast real-time flow-weighted echo-planar imaging (SURF-EPI; **Fig. 1**) that achieves inflow-weighted CSF flow imaging at a high sampling rate (temporal resolution: 46 ms) and an in-plane spatial resolution of 1.3 x 1.3 mm^2^. This high spatiotemporal resolution is achieved by single slice imaging and narrowing the field of view using saturation bands. In addition, the wide dynamic range of SURF-EPI enables simultaneous sensitivity to both neurovascular blood flow and CSF motion. This facilitates interrogation of interactions between venous outflow (internal jugular veins), arterial inflow (internal carotid and vertebral arteries), and CSF dynamics across both time and frequency domains. In particular, arterial signal fluctuations can be used to extract the cerebral arterial input function, allowing reconstruction of CSF cardiac phases without reliance on external gating. As with other inflow-weighted imaging techniques, SURF-EPI captures CSF flow dynamics as changes in voxel signal intensity. We show that the CSF flow waveform derived from SURF-EPI approximates that of cardiac-gated PC MRI, suggesting the potential for calibrating signal intensity to velocity. Thus, our approach complements conventional PC MRI by enabling ultrafast CSF flow imaging independent of velocity encoding, reducing sensitivity to dynamic range limitations and phase noise, while permitting simultaneous assessment of neurovascular and CSF flow dynamics.

**Figure 1.**
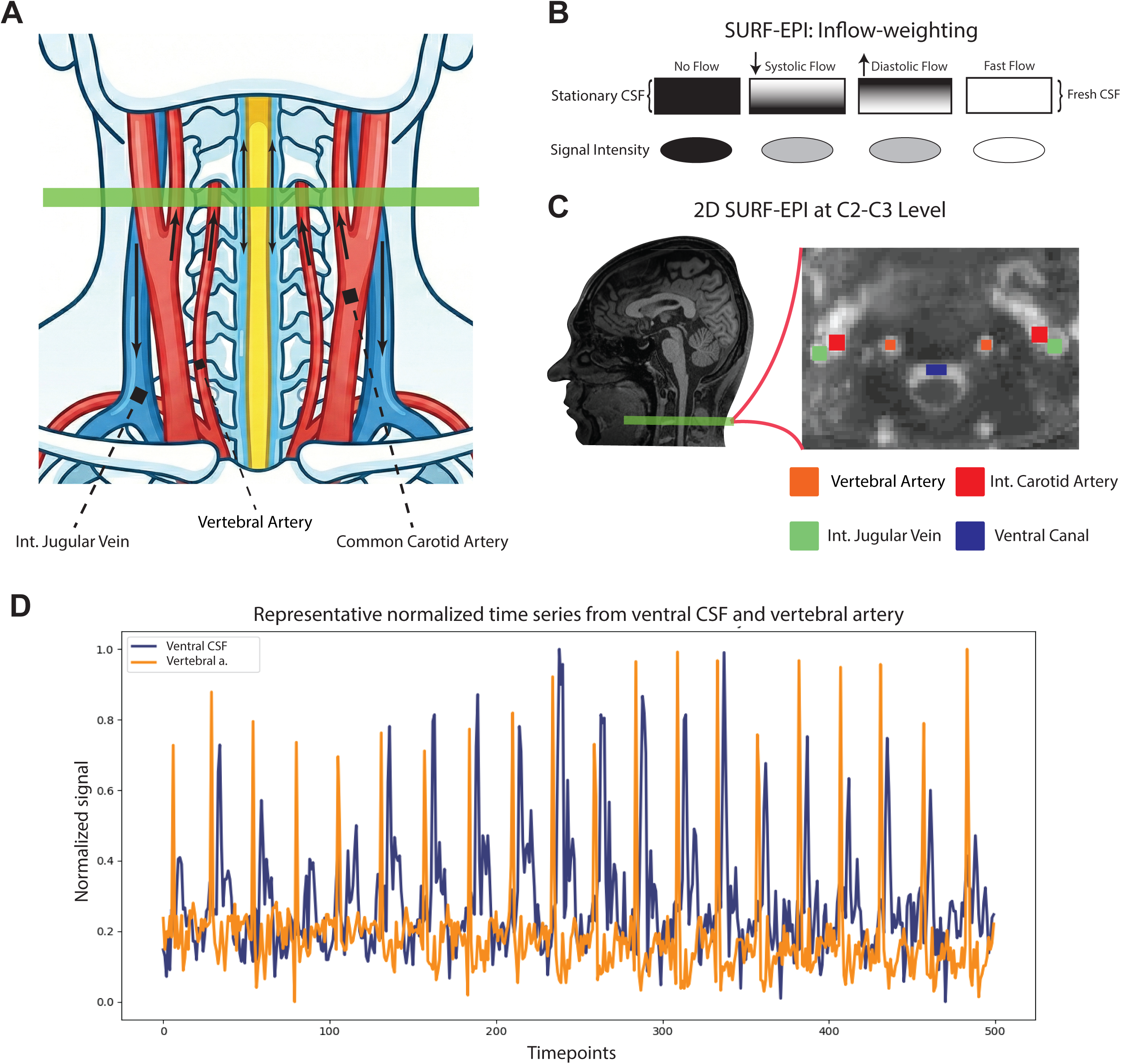
MR physics basis of SURF-EPI and its application for capturing cervical vascular and CSF pulsatility. **(A)** Coronal schematic of cervical neurovascular anatomy at the C2–C3 level, showing the spatial relationship between vertebral arteries, carotid arteries, internal jugular veins, and the spinal canal. The green bar denotes the 2D SURF-EPI slice position. **(B)** MR physics principle underlying SURF-EPI inflow-weighting. Because the sequence uses a very short TR and large flip angle with repeated excitations, stationary water molecules remain saturated and yield near-zero signal. In contrast, through-plane flow brings fresh, unsaturated spins into the imaging slice, producing bright signal. **(C)** Example 2D SURF-EPI scan at C2–C3 showing the temporal standard deviation of the SURF-EPI signal over time (log scale). Regions of interest include the vertebral arteries (orange), internal carotid arteries (red), internal jugular veins (green), and ventral spinal canal CSF (blue). **(D)** Representative normalized time series from ventral CSF and vertebral artery regions of interest acquired with SURF-EPI. Signals were normalized for visualization of temporal features, as arterial pulsation amplitudes substantially exceed those of CSF.

## 2. Methods and Materials

### 2.1. Participants

Twenty-one adult participants (age: 27.8 ± 6.3, 21 – 44 years; 10 female) were recruited from the Washington University in St. Louis community. Exclusion criteria were primary CSF disorders (e.g., idiopathic intracranial hypertension, hydrocephalus, CSF leak), neurodegenerative disorders, major systemic illnesses, uncontrolled diabetes or hypertension, or any contraindication for MRI. All experimental procedures were approved by the Washington University School of Medicine Institutional Review Board. Written informed consent was obtained from all participants, and all participants received financial compensation for participation.

### 2.2. MRI acquisition

All images were acquired using a 3T Siemens Prisma scanner (software version: XA30, Siemens Healthineers, Erlangen, Germany) equipped with a 64-channel head coil. In a subset of participants (n=14), cardiac (photoplethysmography) and respiratory (respiratory belt) signals were continuously acquired at 40 Hz using an MR-compatible physiological monitoring system integrated with the scanner. All relevant code developed for this study is publicly available (https://github.com/BRAFTI/surf-epi).

#### SURF-EPI

Ultrafast single-slice EPI images were acquired with large flip angle and short repetition time (TR) to enhance inflow-weighting^17^ and achieve high temporal resolution (Fig. 1). Anterior and posterior saturation bands were utilized to enable small field-of-view imaging. SURF-EPI acquisitions were performed at the C2–C3 vertebral disc level (Fig. 1), oriented perpendicular to the spinal cord under task-free conditions, allowing free breathing and the natural occurrence of physiological fluctuations without external stimuli or controlled breathing instructions. Acquisition parameters were as follows: TR: 46 ms (sampling rate: 21.7 Hz), TE: 14 ms, flip angle: 90°, slice thickness: 5 mm, in-plane resolution: 1.3 × 1.3 mm^2^, FOV: 125 × 94 mm^2^, and 3,000 time points (141 seconds). Parallel imaging was employed using modified sensitivity encoding (mSENSE)^21^ with an acceleration factor of 2 and phase-encoding partial Fourier of 6/8.

#### Phase-contrast MRI

Phase-contrast MRI was performed at the C2–C3 level using the same slice orientation as SURF-EPI, with a velocity encoding (VENC) of 8 cm/s. Acquisition parameters were as follows: TR: 66.6 ms, TE = 7.21 ms, flip angle = 10°, slice thickness: 5 mm, in-plane resolution: 0.6 × 0.6 mm^2^, FOV: 160 × 160 mm^2^, calculated phases = 40, 2 signal averages, acquisition time: 2:52.

### 2.3. SURF-EPI analysis

For all SURF-EPI analyses, regions of interest (ROIs) were defined using FSLeyes (part of FSL v6.0.7)^22^ to manually delineate key neurovascular and CSF compartments, including the ventral spinal canal (anterior to the spinal cord), dorsal spinal canal (when present), vertebral arteries, internal carotid arteries, and internal jugular veins. Signal processing was performed using SciPy (v1.11.0).

#### Power spectral analysis

Power spectral density (PSD) was computed using the welch function from the *scipy.signal* library, applied to ROI-averaged time series. To account for inter-individual variability in cardiac and respiratory rates, power spectra were rescaled so that the cardiac peak was aligned to a unitless normalized frequency of 1, and the respiratory peak to 0.25. This normalization enabled comparison of physiological signals across participants independent of absolute rate differences in cardiac and respiratory rates. Each spectrum was interpolated using *scipy.interpolate.interp1d* for resampling onto a shared frequency grid. To assess physiological coupling between CSF and vascular signals, magnitude-squared coherence was calculated using the *coherence* function from *scipy.signal*, with nperseg=128, applied to pairs of region-averaged time series. Coherence spectra were rescaled using the same subject-specific frequency alignment and interpolated to a common frequency grid for group-level averaging and analysis.

#### Cardiac cycle detection and reconstruction

Voxelwise arterial waveforms (SURF-EPI signal time series) were extracted from vertebral artery ROIs. These waveforms were standardized by z-scoring. Cardiac systolic peaks were identified using the *find_peaks* function from the *scipy.signal* module with a minimum peak height of 0.5 (arbitrary units) and a minimum inter-peak interval of 670 ms to suppress false detections. The voxel with the greatest number of systolic peaks was selected for further analysis; in cases of ties, preference was given to the voxel with the highest power in the 0.8–1.2 Hz frequency (cardiac) band, ensuring selection of the voxel with the strongest cardiac frequency driven signal. Cardiac cycles were defined as the interval spanning seven time points (∼330 ms) before one vertebral artery systolic peak to seven time points before the next, with the preceding window assumed to capture the CSF diastolic phase. For each voxel, time series segments of average cycle length were extracted, aligned to systolic peaks, and averaged to yield a representative gated waveform.

### 2.4. Cardiac Cycle–Resolved CSF Flow Metrics

Within the ventral and dorsal spinal canal CSF ROIs, the following CSF flow metrics were calculated: (i) CSF *systole/diastole flow ratio (unitless)*, defined as the ratio of the peak systolic signal intensity to the mean signal intensity during late diastole, computed over the final four diastolic time points of the cardiac cycle within the CSF ROI; (ii) *beat-to-beat variability measures* (unitless), computed as the coefficients of variation across cardiac cycles for (a) peak systolic signal intensity, (b) late diastolic signal intensity, and (c) the systole/diastole signal intensity ratio; (iii) *the fraction of extreme cycles*, defined as the proportion in which peak systolic or late-diastolic signal exceeded twice the corresponding amplitude of the ROI-averaged waveform.

### 2.5. Comparison of CSF and neurovascular flow waveforms derived from SURF-EPI and PC-MRI

CSF flow waveforms reconstructed from SURF-EPI data were compared to absolute waveforms extracted from phase-contrast MRI to assess agreement independent of flow direction. In both modalities, mean signal waveforms were extracted from the ventral spinal canal ROI. The SURF-EPI CSF flow waveform was interpolated to match the temporal resolution of the PC-MRI data. Both waveforms were normalized and aligned by shifting PC MRI signal with a constraint that the peak systole in PC-MRI does not precede the arterial peak in SURF-EPI signal, ensuring consistent systolic-phase alignment. Similarity between CSF flow waveforms derived from the two modalities was quantified using the Pearson correlation coefficient.

### 2.6. Statistical analysis

To validate our self-gated cardiac cycle detection, we compared interbeat intervals derived from vertebral artery peak systole time points in SURF-EPI with those obtained from pulse oximeter recordings (index finger) in the physiological log. Agreement between the two methods was assessed using Bland–Altman analysis. The intraclass correlation coefficient (ICC) was additionally calculated to quantify the consistency of the measurements. Paired t-tests were used for pointwise comparisons of ventral versus dorsal CSF flow dynamics, as well as for comparisons between ventral CSF and neurovascular signals in the power spectral density domain across the frequency spectrum (0.02–5 Hz). For all spectral analyses, *p* values were corrected for multiple comparisons across frequencies using the false discovery rate, with *p*_FDR_ < 0.05 considered statistically significant. Extracted parameters of the CSF flow waveform and its variability were compared between ventral and dorsal CSF using paired t-tests. All statistical analyses were performed in R (v4.3.1).

## 3. Results

### 3.1. Spectral signatures reveal distinct flow dynamics in CSF and neurovasculature

For frequency domain analysis, we first examined the power spectra of CSF flow signal within the ventral spinal canal and blood flow signals from the internal jugular vein, vertebral artery, and internal carotid artery at the C2–C3 level. Notably, average cardiac and respiratory rates could be accurately estimated directly from the power spectra without the need for physiological log data (intraclass correlation coefficients:

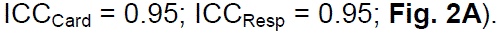

The averaged spectrum revealed prominent peaks at respiratory and cardiac frequencies, along with higher-order cardiac harmonics, in both ventral spinal canal CSF and neurovascular ROIs including the internal jugular veins, vertebral arteries, internal carotid arteries (**Fig. 2B**). Unlike the vascular flow spectra, the ventral CSF exhibited distinctive side peaks flanking the fundamental cardiac frequency. Pointwise comparisons of the power spectrum showed significantly greater power in the ventral CSF at the flanks of the fundamental cardiac frequency compared to all vascular ROIs (*p*_FDR_ < 0.05), suggesting it reflects neurophysiological fluid dynamics unique to the CSF rather than heart rate variability. To determine whether the apparent spectral broadening reflected a true physiological effect rather than an artifact of intersubject averaging, we quantified the full width at half maximum (FWHM) of the cardiac peak in the power spectrum. The FWHM was significantly greater in ventral CSF than in vascular flow spectra (paired t-tests on log-transformed FWHM: vertebral artery: *p* < .001; internal carotid artery: *p* = 0.001; internal jugular vein: *p* = 0.006; **Fig. 2C**). Furthermore, respiratory power in the ventral CSF was significantly higher than in the vertebral (*p*_FDR_ < 0.001) and carotid (*p*_FDR_ = 0.042) arteries, but not the internal jugular vein (*p*_FDR_ = 0.36), indicating that respiration exerts a greater influence on venous and CSF dynamics than on arterial pulsatility (**Fig. 2B**). Conversely, the vertebral artery exhibited higher power at the fundamental cardiac frequency compared to ventral CSF (*p*_FDR_ = 0.013).

**Figure 2.**
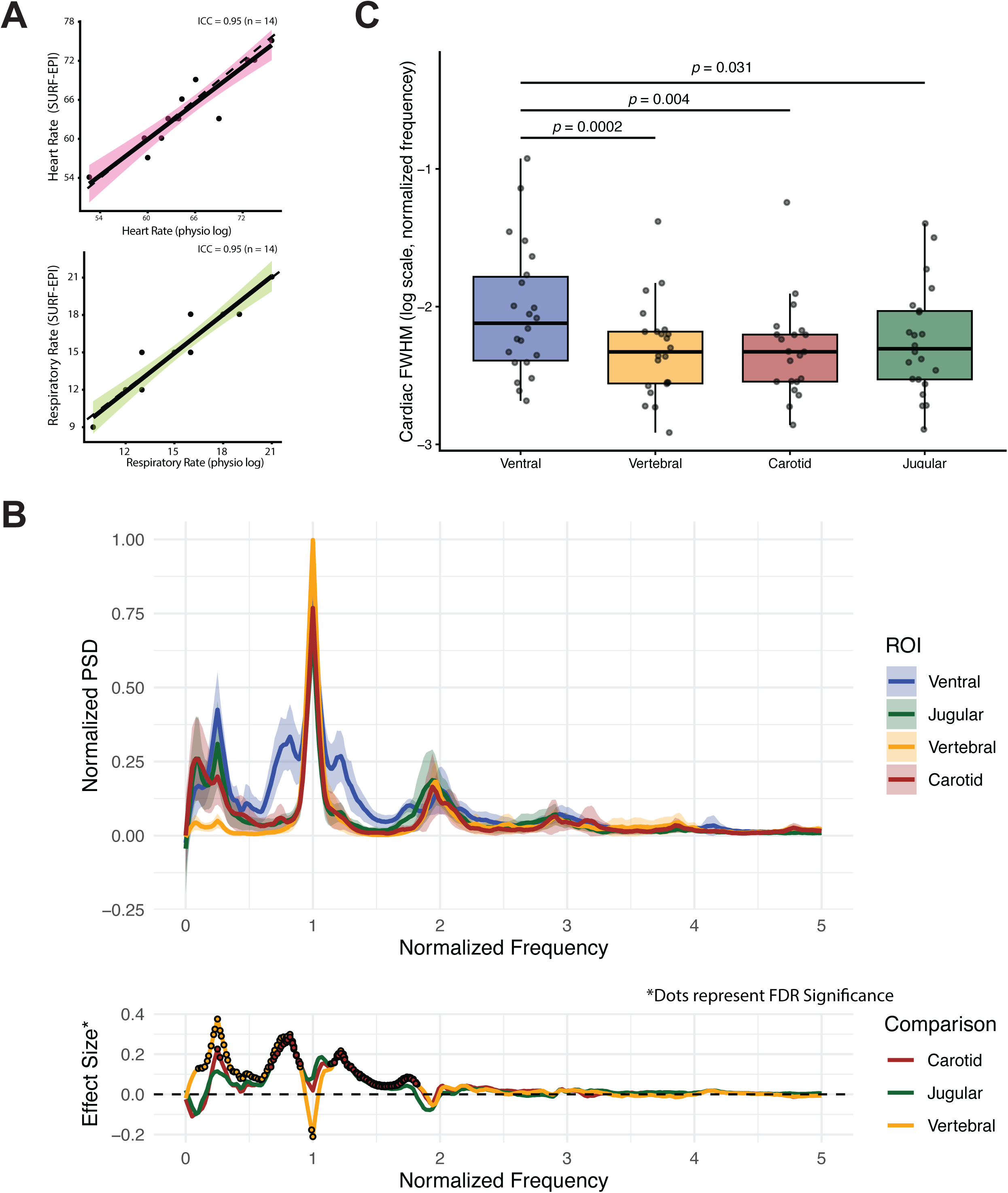
Spectral signatures reveal distinct flow dynamics in CSF and neurovasculature. **(A)** Agreement between physiological monitoring and SURF-EPI-derived estimates of heart rate (top) and respiratory rate (bottom), demonstrating high concordance across participants (intraclass correlation coefficient, ICC = 0.95; *n* = 14). Solid lines indicate linear fits; shaded regions denote 95% confidence intervals. **(B)** Cardiac spectral broadening, quantified as the full width at half maximum (FWHM) of the cardiac peak, across ventral spinal canal CSF and major cervical vessels (y-axis is log-scale). CSF exhibits broader cardiac spectral peaks compared with arterial/venous flow signals, consistent with greater temporal dispersion. **(C)** Group-averaged normalized power spectral density (PSD) of SURF-EPI signals from ventral CSF, internal jugular vein, internal carotid artery, and vertebral artery (top), demonstrating distinct frequency content across fluid compartments. Frequencies are rescaled to cardiac = 1 and respiratory = 0.25. Shaded bands indicate ± standard error. Bottom, frequency-resolved effect sizes relative to ventral CSF, highlighting compartment-specific differences at cardiac and harmonic frequencies; dots indicate frequencies surviving false discovery rate (FDR) correction.

In a subset of participants (n=18) in whom the spinal cord was not abutting the dorsal spinal canal wall, allowing a discernible dorsal CSF column, spectral comparisons were conducted between ventral and dorsal CSF (**Fig. 3A**). Dorsal CSF exhibited lower power at the respiratory frequency (*p*_FDR_ = 0.04) and in the flanks of the cardiac fundamental frequency (low frequency flank: *p*_FDR_ = 0.017; high frequency flank: *p*_FDR_ = 0.029) and the lower frequency flank of its second harmonic (*p*_FDR_ = 0.019), indicating lower respiratory modulation. Conversely, the dorsal column exhibited greater power at the cardiac fundamental frequency (*p*_FDR_ = 0.04).

**Figure 3.**
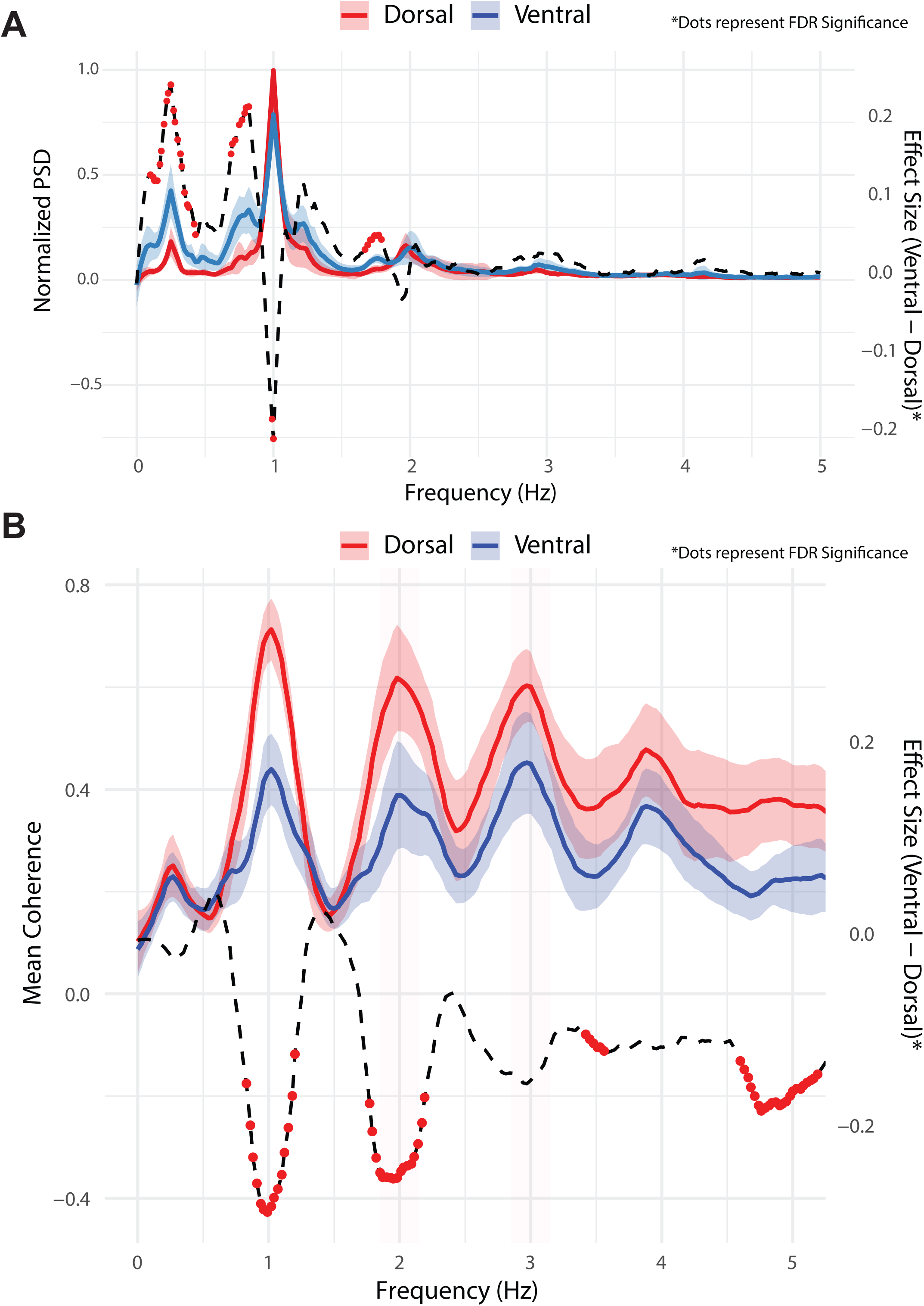
Frequency-dependent differences between ventral and dorsal spinal CSF flow dynamics. **(A)** Group-averaged normalized power spectral density (PSD) of SURF-EPI signals from ventral and dorsal spinal canal CSF columns (n = 18). Ventral and dorsal spinal CSF columns exhibit distinct spectral profiles, with higher power at the fundamental cardiac frequency in dorsal CSF, and relatively greater power at the cardiac flanks and respiratory frequency in ventral CSF. Shaded bands indicate ± standard error. The dashed black trace denotes frequency-resolved effect size (ventral − dorsal); dots mark frequencies surviving false discovery rate (FDR) correction. **(B)** Mean magnitude-squared coherence between CSF and vascular signal in ventral and dorsal CSF as a function of frequency, revealing frequency-dependent coupling between spinal CSF compartments. Shaded bands indicate ± standard error. The dashed black trace represents the frequency-resolved effect size (ventral − dorsal), with dots indicating frequencies surviving FDR correction. The frequency axis is normalized to respiratory = 0.25 and cardiac = 1.

### 3.2. Neurovascular and spinal CSF flow dynamics preferentially couple at cardiac frequencies

We then examined the influence of jugular, vertebral, and carotid signals on CSF flow using coherence analysis. Across all vascular signals, coherence with ventral and dorsal spinal canal CSF flow signal was consistently higher at the fundamental cardiac frequency (0.43 ± 0.17) and its harmonics (second harmonic: 0.39 ± 0.25; third harmonic: 0.45 ± 0.23) compared to the respiratory frequency (coherence: 0.22 ± 0.11), indicating that cardiac pulsations are the primary drivers of vascular-CSF coupling (*p* < 0.001 for all comparisons). Compared to ventral CSF column, the dorsal CSF column exhibited consistently higher coherence with all vascular signals at the fundamental cardiac frequency and its harmonics (cardiac: 0.70 ± 0.12; first harmonic: 0.61 ± 0.21; second harmonic: 0.60 ± 0.14; *p*_FDR_ < 0.05; **Fig. 3B**).

### 3.3. Agreement between SURF-EPI Cardiac Cycle Detection and Physiological Log Data

We next sought to delineate the relationship between the arterial input function and the CSF flow waveform. Cardiac cycles were identified by applying a peak detection algorithm to the vertebral artery time series (**Fig. 4A, 4B**). To evaluate the concordance between cardiac cycle detection from SURF-EPI and physiological recordings, we calculated the ICC between beat-to-beat cardiac cycle lengths derived from each method (**Fig. 4C, 4D**). Mean and proportional bias were negligible, confirming the absence of systematic or magnitude-dependent differences between methods. Among the fourteen participants with simultaneous physiological recordings, 78.6% showed either very strong (ICC > 0.80; 21.4%) or strong agreement (ICC = 0.60–0.79; 57.1%). Moderate agreement (ICC = 0.40–0.59) was observed in 14.3% of participants, and one subject (7.1%) demonstrated only fair agreement, with an ICC of 0.30. Further analysis demonstrated that lower heart rate variability was associated with reduced ICC values (Spearman ρ = 0.51; *p_one-tailed_ =* 0.03). These results highlight the robustness of SURF-EPI for concurrent assessment of CSF and vascular dynamics, with cardiac cycle detection closely matching physiologic recordings in most participants. Importantly, vertebral artery waveforms derived from SURF-EPI may provide an advantage over conventional peripheral pulse recordings by capturing the arterial input to the brain with greater anatomical specificity.

**Figure 4.**
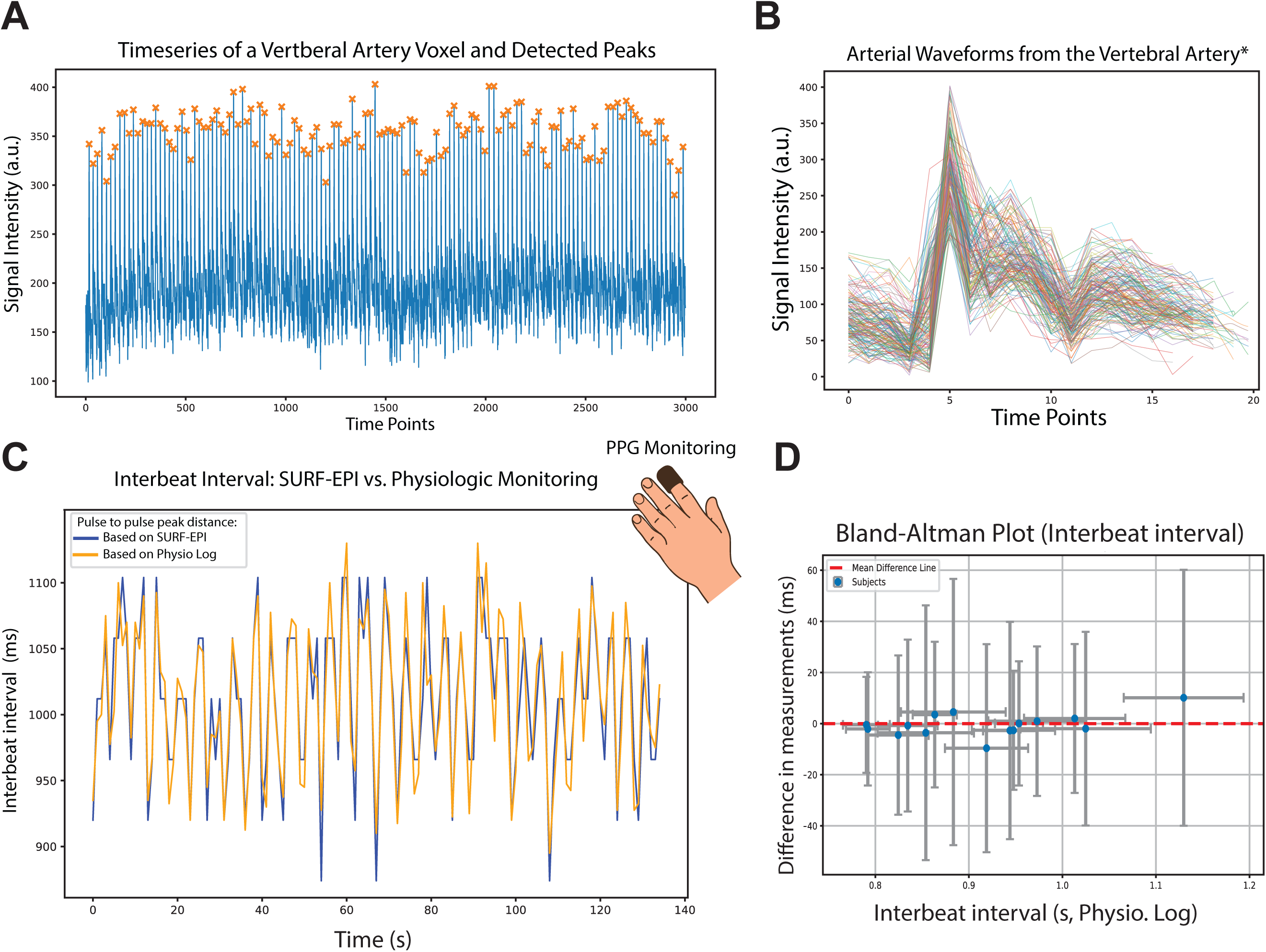
Vascular signal time series enable self-gated cardiac cycle detection in SURF-EPI. **(A)** Representative vertebral artery time series with detected systolic peaks used for cardiac cycle identification. **(B)** Arterial waveforms extracted from the vertebral artery and aligned to detected systolic peaks, illustrating pulse morphology across cycles. **(C)** Example subject showing beat-to-beat interbeat intervals derived from SURF-EPI and physiological pulse recordings. **(D)** Bland–Altman analysis of interbeat intervals across subjects. Each point represents one subject, with the x-axis showing the mean cardiac cycle length and the y-axis showing the mean difference between SURF-EPI–derived and physiologic interbeat intervals. Vertical and horizontal error bars indicate the standard deviation of interbeat interval differences and cycle lengths, respectively.

### 3.4. Alignment of CSF flow waveforms between SURF-EPI and PC-MRI

CSF flow waveforms were extracted from a ventral spinal canal ROI at the C2–C3 level in both SURF-EPI and PC MRI datasets (**Fig. S1**). To enable direct comparison, SURF-EPI signals were up-sampled to match the number of cardiac phases in the phase-contrast MRI, and PC MRI waveforms were converted to absolute values to account for the non-directionality of SURF-EPI. Waveform similarity was assessed using correlation analysis. In 76.2% of participants, either very strong (r ≥ 0.80; n = 4; 19.0%) or strong (r = 0.60–0.80; n = 12; 57.1%) correlation was observed between waveforms from the two modalities. The remaining five participants (23.8%) demonstrated moderate correlation (r = 0.52–0.59). Collectively, these findings suggest that SURF-EPI–derived CSF flow waveforms exhibit strong concordance with phase-contrast MRI, with the highest agreement observed near CSF systole.

### 3.5. Dorsal spinal CSF flow demonstrates lower cycle-to-cycle variability

SURF-EPI not only reproduced the average CSF waveform across the cardiac cycle (analogous to phase-contrast MRI) but also enabled quantification of beat-to-beat variability. To compare ventral and dorsal CSF columns at C2–C3, we assessed the systole/diastole flow-related signal ratio (peak systolic signal divided by mean late-diastolic signal), the coefficients of variation across cycles for peak systole, late diastole, and the systole/diastole ratio, and the fraction of cycles in which peak systole or late-diastolic signal exceeded twice their respective means (**Fig. 5**). Ventral CSF exhibited significantly greater variability across all three measures (peak systole, *p* = 1.2 × 10^−4^; late diastole, *p* = 0.0035; systole/diastole ratio, *p* = 0.0018) and a higher proportion of extreme systolic peaks (*p* = 0.047). There was also a trend toward a higher mean systole/diastole ratio (p = 0.058) and more frequent extreme late-diastolic fluctuations (p = 0.093) in the ventral spinal canal. Together, these findings show that ventral CSF flow exhibits greater cycle-to-cycle variability.

**Figure 5.**
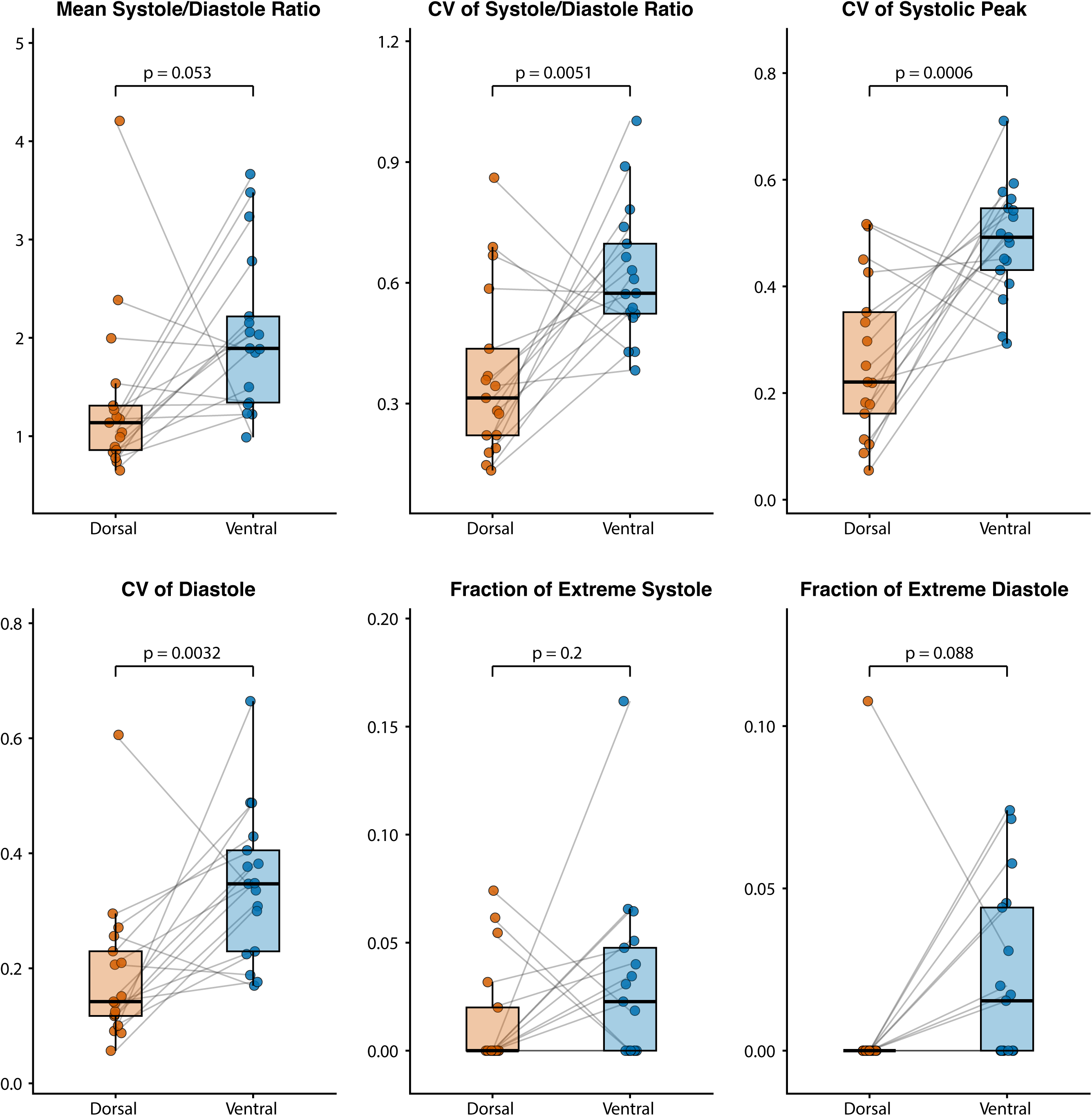
Ventral–dorsal differences in cardiac-cycle variability and systolic–diastolic flow dynamics. Metrics included the mean systole-to-diastole flow ratio (defined as peak systolic signal divided by mean late-diastolic signal), the coefficient of variation (CV) of this ratio across cardiac cycles, the CV of peak systolic and late-diastolic signals, and the fraction of cardiac cycles exhibiting extreme systolic or late-diastolic fluctuations (signal > 2× subject mean).

## 4. Discussion

Our findings establish SURF-EPI as a simple, robust method for capturing real-time CSF and vascular dynamics simultaneously without the constraints of velocity encoding or external gating. While the average CSF waveform derived from SURF-EPI generally paralleled phase-contrast MRI, the method also enabled spectral decomposition and cycle-to-cycle variability analysis that extend beyond what conventional PC MRI provides. In this sense, SURF-EPI offers not only an acquisition strategy but also an analytic framework for probing CSF flow in both the time and frequency domains.

In the ventral CSF, we observed spectral widening around the fundamental cardiac frequency, a feature that was absent in vascular signals. This broadening may reflect the viscoelastic and compliance properties of the intracranial system, which is in communication with the ventral spinal subarachnoid space^23,24^. Prior studies suggest that the cranium normally functions as a pulsation absorber, acting like a notch filter that attenuates the fundamental cardiac frequency^25^. Another explanation is that these represent sidebands generated by respiratory modulation of the cardiac pulsation, a phenomenon recently characterized with magnetic resonance encephalography (MREG) as cardiorespiratory envelope modulation^26,27^. In addition, respiratory contributions were more prominent in the ventral CSF and internal jugular veins than in the carotid or vertebral arteries, consistent with the dominant influence of respiration on venous return and CSF flow dynamics^12,18,28^.

Compared with the dorsal spinal CSF column, the ventral canal exhibited a more complex spectral profile, with greater respiratory and non-cardiac contributions, higher beat-to-beat variability, and lower coupling with neurovascular flow. This divergence likely reflects anatomical and functional differences between the compartments, as the spinal subarachnoid space is structurally compartmentalized by arachnoid membranes anchored to the denticulate ligaments and nerve rootlets, resulting in distinct ventral and dorsal cisternal spaces^29,30^. Furthermore, the ventral CSF space serves as the primary conduit between the spinal canal and the cranial vault^31^. Consistent with this, time-spatial labeling inversion pulse (time-SLIP) imaging has shown that respiration drives substantial bidirectional motion in the prepontine cistern, with cephalad displacement during inspiration and caudad displacement during expiration^32^. Additional factors may contribute to the observed compartmental differences, including posture-dependent displacement of the spinal cord within the thecal sac during supine imaging, which effectively enlarges the ventral subarachnoid space, and differences in vascular anatomy, with a single anterior spinal artery ventrally and paired posterior spinal arteries dorsolaterally, potentially shaping local pulsatility and CSF–vascular coupling.

Spectral analysis has been used with inflow-weighted imaging to probe low-frequency CSF flow fluctuations^16,19^. Yet, limited sampling rates (2–5 Hz) constrained prior studies to slow oscillatory components. SURF-EPI, with effective sampling rates exceeding 20 Hz, resolves higher-frequency content, including the cardiac fundamental and its harmonics. Related work with MREG has demonstrated whole-brain sampling at ∼10 Hz by employing strong k-space undersampling^27^. However, MREG is fundamentally a T2*-weighted blood oxygen level dependent (BOLD) technique, yielding a complex composite of vascular, brain tissue, and CSF signals at relatively low spatial resolution. Thus, while valuable for mapping global brain pulsations, MREG cannot isolate CSF flow from vascular dynamics. At the highest temporal resolution, pencil-beam imaging samples up to ∼40 Hz, yet its one-dimensional acquisition sacrifices spatial mapping and cannot simultaneously assess blood and CSF signals^33,34^.

Rapid imaging of neck arteries permitted use of their flow waveform as an arterial input function for self-gating. Physiological gating methods introduce additional complications. In PC MRI, cardiac gating often relies on fingertip photoplethysmography, which is subject to variable time lags (pulse arrival time) influenced by factors such as hypertension, owing to the delay between cardiac contraction and peripheral pulse detection^35^. While EKG gating offers more direct cardiac timing and minimizes temporal variability in CSF flow measurements, its practical challenges (e.g., the need for chest shaving to ensure reliable skin contact) often restrict its use to smaller studies^36,37^. By contrast, simultaneous imaging of neurovascular and CSF flow enables precise temporal characterization of CSF dynamics with respect to the cerebral arterial input function, avoiding the limitations of peripheral gating.

The mean CSF flow waveform derived from SURF-EPI showed strong correlation with PC MRI across the cardiac cycle in most subjects, although in some individuals the correlation was only moderate. This discrepancy likely reflects a fundamental difference in flow weighting: PC MRI encodes directionality, such that bidirectional flow within a cardiac phase can cancel out, yielding a net signal near zero. In contrast, SURF-EPI is sensitive to absolute flow regardless of direction, meaning that phases with opposing bidirectional flow contribute high signal intensity rather than nulling. Consistent with recent real-time PC MRI studies, respiratory influences are most evident during late diastole, when cranio-caudal reversals are frequent, whereas early systolic peaks—dominated by caudal flow—remain comparatively unaffected^14^. Taken together, SURF-EPI complements PC MRI by providing high-temporal-resolution signatures of CSF motion, while PC MRI supplies directional velocity information. When correlation is high and flow direction is more certain (e.g., early systole), PC MRI could in principle be used to calibrate SURF-EPI signal intensity to velocity units.

Compared with real-time PC MRI, which provides quantitative velocity and directionality, the key advantages of SURF-EPI are its wide dynamic range and temporal fidelity. In PC MRI, the selection of a single VENC imposes a narrow dynamic range, precluding simultaneous visualization of high-velocity blood flow (20–120 cm/s) and low-velocity CSF flow (5–15 cm/s) within the same sequence. By contrast, the sensitivity of SURF-EPI to a broad range of flow velocities enables concurrent characterization of CSF and vascular flow patterns, allowing assessment of CSF–blood coupling. Beat-to-beat analysis further showed that CSF flow can exceed twice the average peak amplitude in up to 12% of cycles, fluctuations that would introduce aliasing and measurement errors in PC MRI but are captured by SURF-EPI. Given that each SURF-EPI frame is fully sampled, rather than reconstructed from k-space sub-sampling with k-space-time acceleration^7,38,39^, the method achieves true high temporal resolution without temporal blurring, while maintaining high in-plane spatial resolution. However, SURF-EPI lacks explicit velocity encoding, making it best viewed as complementary to conventional PC MRI. Used together, the two techniques can separate the role of real-time dynamic characterization from quantitative velocity measurement, with the potential for SURF-EPI signals to be calibrated against PC MRI to provide an integrated framework for neurofluid imaging.

Several limitations warrant consideration. First, our analyses were rescaled to cardiac and respiratory frequencies, and thus other sources of very low-frequency variance may not have been fully captured, as they are not necessarily coupled to these frequencies. Second, CSF dynamics were quantified using ventral and dorsal subarachnoid space ROIs, which may mask finer-scale spatial heterogeneity. Future voxel-wise approaches could resolve spatiotemporal CSF flow patterns beyond ROI-level analyses. Third, the present implementation of SURF-EPI was limited to single-slice acquisitions; however, simultaneous multislice imaging could enable concurrent assessment of spinal and intracranial CSF flow without significant time penalty^40^. Fourth, our study was restricted to healthy subjects, and future investigations in patient populations will be essential to determine how alterations in CSF dynamics manifest across time and frequency domains and how they relate to intracranial pathophysiology, including structural abnormalities, disturbed CSF circulation, and altered viscoelastic properties. Finally, although a frame rate of 20 Hz exceeds the Nyquist limit even for cardiac harmonics, precise evaluation of phase shifts and characterization of vascular and CSF waveforms benefit from even higher temporal resolution. To this end, advanced undersampling and acceleration strategies could be leveraged to achieve higher sampling rates^41^.

In conclusion, SURF-EPI provides a simple and robust means to capture real-time CSF and neurovascular dynamics with high temporal fidelity and a wide dynamic range. By enabling simultaneous assessment of arterial inflow, venous outflow, and CSF motion, SURF-EPI reveals spectral and cardiac cycle-resolved features that conventional PC-MRI cannot capture. More broadly, the correspondence between SURF-EPI and PC-MRI waveforms suggests that quantitative velocity measurements from PC-MRI could be combined with the fast, non-directional dynamics captured by SURF-EPI to yield unified, calibratable markers of CSF flow in real time. Beyond cardiac-cycle–locked dynamics, the real-time nature of SURF-EPI provides a framework for probing off–cardiac-cycle CSF flow patterns and task-mediated perturbations, such as Valsalva maneuvers or coughing, enabling interrogation of transient, physiologically evoked CSF responses. By providing a simple, robust, and broadly deployable method for real-time CSF imaging, SURF-EPI enables investigation of CSF flow dynamics across health and disease and lays a foundation for future applications spanning intracranial pressure–volume disorders, hydrocephalus, CSF flow obstruction, and neurovascular disorders.

## Data and Code Availability

All relevant code developed for this study is publicly available (https://github.com/BRAFTI/surf-epi). Imaging data supporting the findings of this study are available from the corresponding author upon reasonable request.

## Author Contributions

A.H.S. and A.N. conceived and designed the study; A.H.S. and A.N. developed the methodology; A.H.S., C.E., H.H., and A.N. acquired the data; A.H.S., A.H.G.A., and A.N. performed the formal analysis; A.N. and A.H.S. wrote the original draft; and all authors (A.H.S., A.H.G.A., B.A.N., A.S., C.E., H.H., J.S.S., D.D.L., J.M.S., and A.N.) reviewed and edited the manuscript. A.N. supervised the study.

---

**Figure S1.** (A) Phase-contrast (PC) MRI slice at the C2–C3 level used for comparison with SURF-EPI, with the ventral spinal canal ROI indicated. (B) CSF flow waveforms from all participants exhibiting very strong agreement between SURF-EPI and PC MRI after temporal alignment and normalization. (C) CSF flow waveforms from all participants exhibiting strong agreement between SURF-EPI and PC MRI. (D) CSF flow waveforms from all participants exhibiting moderate agreement between SURF-EPI and PC MRI.

