## Supplementary figures and images for "Simultaneous Real-time Imaging of Neurofluid and Neurovascular Dynamics Using Ultrafast Flow-weighted Echo-Planar Imaging"

### Supplementary Figure S1

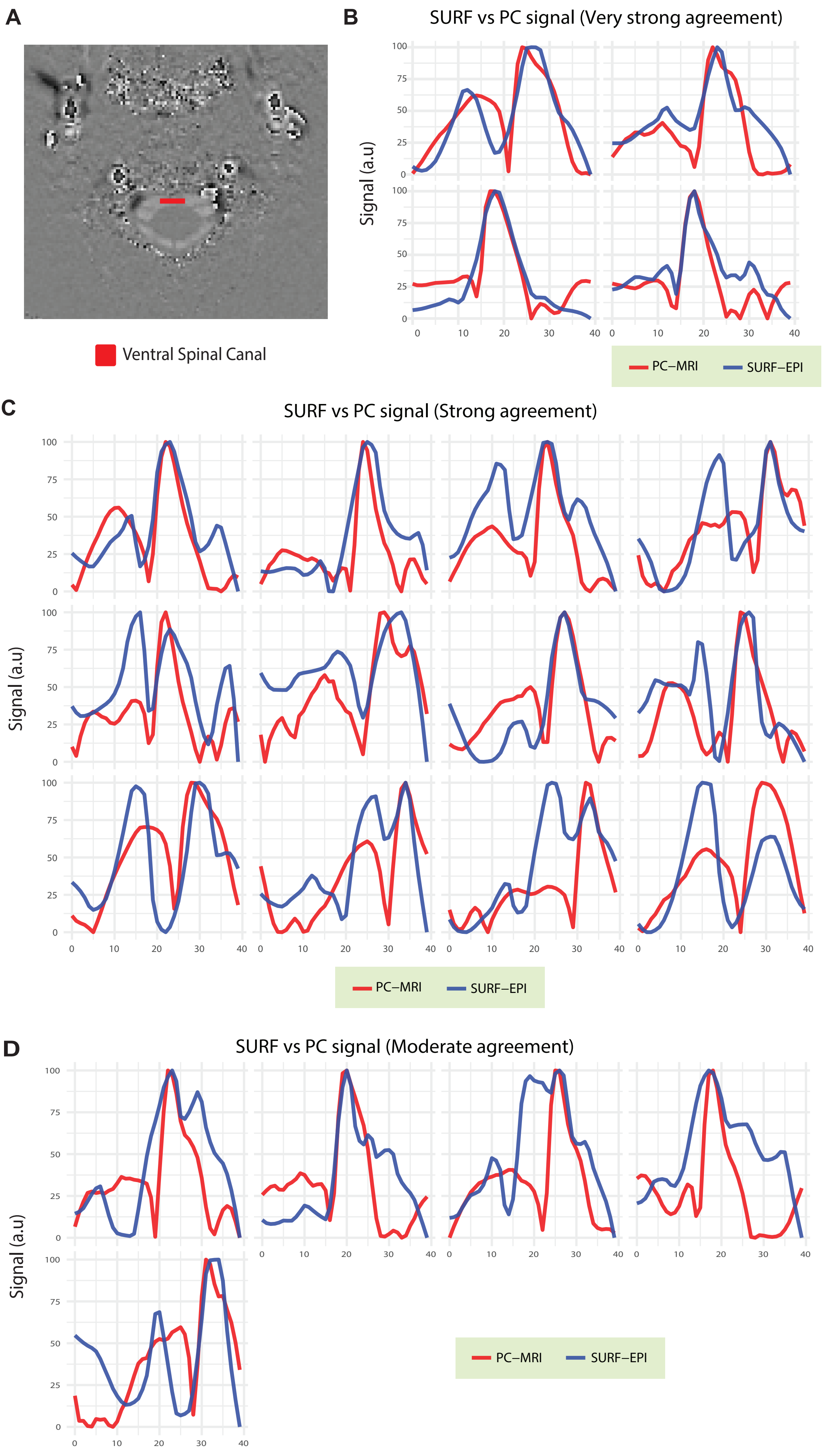
